## Supplementary figures and images for "Extensive diversity in *Escherichia coli* Group 3 capsules is driven by recombination and plasmid transfer from multiple species"

### Supplemental Fig S1

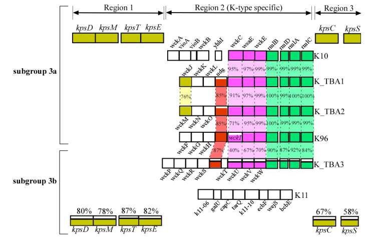
